# Temporal landscape and translational regulation of A-to-I editing in mouse retina development

**DOI:** 10.1101/2023.03.07.531644

**Authors:** Ludong Yang, Liang Yi, Jiaqi Yang, Rui Zhang, Zhi Xie, Hongwei Wang

## Abstract

The significance of A-to-I RNA editing in neurological development is widely recognized, however, its influence on retina development remains to be thoroughly understood. In this study, we aimed to characterize the temporal landscape of A-to-I editing in mouse retina development using total RNA-seq and Ribosome profiling, with a specific emphasis on its effect on gene translation. Our findings revealed that the editing underwent plastic changes and distinct editing contents facilitated stage-specific functions. Further analysis showed a dynamic interplay between A-to-I editing and alternative splicing, with A-to-I editing having a significant impact on splicing efficiency and increasing the quantity of splicing events. A-to-I editing held the potential of enhancing the translatome’s diversity, but this came at the expense of reduced translational efficiency. When coupled with splicing, it could produce a coordinated regulatory effect on translatome dynamics. Overall, our study presents a temporally resolved atlas of A-to-I editing, connecting its dynamic changes with the regulatory impact on splicing and translation.

## INTRODUCTION

Adenosine to inosine (A-to-I) RNA editing, catalyzed by the ADAR (adenosine deaminases acting on RNA) family of enzymes, is the most prevalent type of RNA editing in metazoan. This process transforms adenosine into inosine in double-stranded RNA^1–3^. As a result of this editing, the ribosomes and splicing machinery interpret inosines as guanosines, leading to potentially non-synonymous substitutions if the editing occurs within the coding part of a transcript, thereby producing novel protein variants^4,5^. Thus, A-to-I editing has a profound impact on targeted RNAs, not only altering their sequences compared to the genome but also influencing the cellular destiny of RNA molecules. To date, screening of the editome has revealed numerous A-to-I editing sites in various primate species^6,7^. There is growing evidence that A-to-I editing plays a crucial role in neurological development, performing various physiological functions such as regulating neuronal transmission, modulating synaptic plasticity, and controlling the timing of neurogenesis^8–11^. Nevertheless, given the increasing identification of A-to-I editing sites, the majority of them still have undefined spatial, temporal, and functional characteristics.

The survey of A-to-I editing in mammalian development not only unveils dynamic nature of RNA editing, but also sheds light on the intricate regulation of this process. The brain, being the primary focus of developmental editing due to its high prevalence of A-to-I editing in the central nervous system (CNS) of mammals, is the target of many studies^12^. However, the retina, as an extension of the CNS, is also a valuable organ to study neurodevelopment due to its accessibility. Despite this, the A-to-I editing in the developing retina is poorly understood, with only one prior study investigating this phenomenon. This existing study was limited in scope, only providing a snapshot of editing at 2,000 shared sites^13^, making it challenging to fully comprehend the prevalence, consequences, and significance of A-to-I editing in the retina.

A-to-I editing represents a fascinating layer of gene expression regulation that has been recognized for its role in illuminating the molecular basis of retina development. Numerous studies have shown that A-to-I editing plays a dynamic role in regulating transcriptome diversity and fine-tuning gene expression, including through interactions with alternative splicing^14–16^. However, the impact of A-to-I editing on translatome diversity and translational regulation during retina development remains an underexplored area. This is especially noteworthy since the central dogma of gene expression involves two sequential steps: transcription and translation.

Herein, we focused on exploring the temporal landscape and translational regulation of A-to-I editing during the development of mouse retina. To achieve this, we employed ultradeep transcriptomic data to evaluate the A-to-I editing profiles during three crucial stages of the visual system’s development: embryonic, neonatal and eye-opening. We then analyzed the temporal dynamics of A-to-I editing during development, observing distinct editing contents that facilitated stage-specific functions, as well as temporal editing patterns that demonstrated the editing landscape’s plasticity in response to development. Additionally, we investigated the impact of A-to-I editing on alternative splicing and found that A-to-I editing precedes splicing and has a widespread effect on splicing. Finally, we performed ribosome profiling (Ribo-seq) to examine the regulatory effect of A-to-I editing to gene translation, revealing that it acts as a buffer to diminish the efficiency of translation and produces coordinated effects when coupled with splicing. In conclusion, our study highlights the crucial role that A-to-I editing plays in regulating temporal patterning and functional specification in the developing mouse retina.

## MATERIALS AND METHODS

### Tissue collection

Specimens of retinal tissue were obtained from C57BL/6 mice of wild-type origin, which were supplied by the Animal Centre of Southern Medical University in Guangzhou, China. The tissue samples covered a diverse spectrum of retina development phases, encompassing embryonic (13.5 days), neonatal (postnatal 0 and 6 days), and eye-opening (postnatal 21 and 42 days) stages. After crushing in a tissue mashing machine, the samples were immediately frozen in liquid nitrogen to preserve their quality. All animal experiments were approved by the Animal Ethics Committee of the Zhongshan Ophthalmic Center, Sun Yat-sen University (Guangzhou, China) with the permit number 2017-085A.

### Library preparation and sequencing

The total RNA-seq and Ribo-seq libraries for each sample were generated according to previously reported protocols^17^. In brief, the tissue samples were lysed using a mixture of mammalian lysis buffer (200 μl 5x Mammalian Polysome Buffer, 100 μl 10% Triton X-100, 10 μl DTT (100 mM), 10 μl DNase I (1U/μl), 2 μl Cycloheximide (50 mg/ml), 10 μl 10% NP-40, and 668 μl Nuclease-Free Water). After 20 minutes of incubation on ice, the lysates were clarified through centrifugation at 10,000xg for 3 minutes at 4°C. The 300 μl of clarified lysates were then treated with 5 Units of ARTseq Nuclease for 45 minutes at ambient temperature to perform nuclease digestion. The ribosome-protected fragments were purified using Sephacryl S-400 HR spin columns (GE Healthcare Life Sciences) and RNA Clean & Concentrator-25 kit (Zymo). The ribosomal RNA was removed from the purified RNAs using the Ribo-Zero magnetic kit (Epicentre). The Ribo-seq library was constructed using the ARTseq™ Ribosome Profiling Kit (Epicentre). The 100 μl of clarified lysates were used for total RNA extraction and purification. The purified RNAs were linked to a 5’ adaptor, followed by reverse transcription and PCR amplification, culminating in a strand-specific total RNA-Seq library created using the VAHTSTM total RNA-Seq v2 Library Prep Kit from Illumina (Vazyme Biotech). The resulting libraries were then sequenced using the Illumina HiSeq 2500 following the manufacturer’s protocol, producing 2×125 bp paired-end reads for total RNA-seq and 1×51 bp single-end read runs for Ribo-seq.

### Data preprocessing

The raw read data of total RNA-Seq and Ribo-seq were demultiplexed using CASAVA (v1.8.2), and then the 3’-end adapter was removed using Cutadapt (v1.8.1)^18^. To improve the quality of the data, low-quality sequences were trimmed using fastp (v0.20.1)^19^. For the Ribo-seq data, an additional step was taken to filter the reads to retain only those with lengths between 25 and 35. To further clean the data, reads mapped to mouse rRNA and tRNA sequences were excluded. The remaining reads were then realigned to the mouse reference genome (GENCODE, GRCm38.p6) using Hisat2 (v2.1.0)^20^. Only the reads that were uniquely mapped reads were included in the downstream analysis. Finally, the quantity of reads for each gene was calculated using featureCounts (v1.6.4)^21^.

### Identification and annotation of A-to-I editing sites

Picard’s MarkDuplicates (v2.23.3) [https://github.com/broadinstitute/picard] was used to reduce the PCR bias on the editing level. RNA editing sites were then detected using REDItools2^22^ with a parameter of -s 2. To ensure the accuracy of the editing sites, several steps were taken to minimize the risk of false positives, including: (1) preserving sites that existed in both replicates, (2) trimming 12 nucleotides from the start and 2 nucleotides from the end of the reads, (3) eliminating sites located in SNPs (https://hgdownload.soe.ucsc.edu/goldenPath/mm10/database/snp142.txt.gz) and 4 nucleotides of the splicing site, (4) removing sites with multiple variant types, and (5) retaining sites with a minimum editing level of 0.02, at least 10 high-quality reads covered and at least 3 reads edited. The genomic features and amino acid changes of RNA editing sites were annotated using ANNOVAR (2020-06-07 release)^23^. Any sites with ambiguous annotation were excluded. The sequence context surrounding the RNA editing sites was analyzed using motifStack (v1.30.0)^24^. motifStack was utilized to analyze the sequence context surrounding the RNA editing sites. The above identification and annotation of A-to-I editing sites were integrated into a tailored workflow using Snakemake (v6.0.2)^25^.

### Characterization of editing pattern

Mfuzz (v2.46.0)^26^ was used to study the editing patterns of sites during retina development. The following steps were taken: (1) a matrix s created that contained information on the editing level for each editing site in each stage, in order to create an ExpressionSet object, which is required by mfuzz, (2) mfuzz::mestimate was used to determine the optimal fuzzifier value (m), (3) the mfuzz function was employed with parameters: c = 6 and m, determined in the previous step, to cluster editing sites, and finally, (4) the order of editing patterns was reordered based on the their peak timing during development.

### Estimation of gene expression and translational efficiency

The counts of all CDS from the 10 total RNA-seq and 10 Ribo-seq samples were consolidated into a single table, and genes with a total count of less than 10 across all samples were discarded. The DESeq2 (v1.26.0)^27^ tool was used to estimate and remove the library size effect, and the gene expression was corrected based on their lengths to obtain the normalized gene expression. Finally, the translational efficiency was calculated by taking the ratio of the normalized gene expression at the translational level to the expression at the transcriptional level.

### Detection of differential editing sites

The REDIT-LLR function from the REDITs^28^ software was used to identify differentially edited sites between adjacent developmental stages (P0 vs. E13.5, P6 vs. P0, P21 vs. P6, and P42 vs. P21). This was achieved by inputting a 2-row matrix that contained the number of edited and non-edited reads obtained from the output of REDItools2. Only sites with a p-value of 0.05 were considered statistically significant.

### Detection of differential translational efficiency

The DESeqDataSetFromMatrix function was used to identify differentially translational efficiency (dTE) genes between adjacent developmental stages, with a design formula of ‘library type + time + library type:time’. The input for this function was the combined counts obtained from the ‘Estimation of gene expression and translational efficiency’ section. The ‘results’ function was then used to extract dTE genes. Only genes with an absolute log2(fold change) of at least 1 and an adjusted p-value of 0.05, calculated using the Benjamini-Hochberg method, were considered statistically significant.

### Identification of alternative splicing events

Vast-tools (v2.5.1)^29^ was used to identify alternative splicing (AS) events from total RNA-seq alignment BAM files. The parameters --min_Fr 0.2 --noVLOM --p_IR were applied to filter out events that did not have sufficient coverage (at least 25 reads for IR and 15 for others) in at least 20% of the samples and those that did not pass the binomial test. Notably, five distinct types of splicing events were revealed in this analysis, including exon skipping and mutually exclusive exons (EX), intron-retained (IR), alternative acceptors (Alt3), alternative donors (Alt5), and exon skipping for micro exons (MIC).

### Detection of differential splicing events

The ‘compare’ function of vast-tools was used to determine differential splicing events such as the percentage splicing in (PSI) for AS events or percentage intron retained (PIR) for IR events, between adjacent developmental stages.The parameters: --min_dPSI 15, --paired --min_range 5, --noVLOW --p_IR, and -sp mm10 were employed.

### Correlation analysis between RNA editing and splicing

To explore the relationship between RNA editing and splicing, each editing site was linked to its corresponding closed splicing event, excluding intergenic sites that were located outside of gene regions. Pearson’s correlation coefficient was then calculated for each pairing of editing site and splicing event. Based on the correlation strength, the pairings were categorized into four groups: Strong (absolute value (r) ≥ 0.7), Moderate (0.7 > r ≥ 0.5), Weak (0.5 > r ≥ 0.3), and None (r < 0.3). Only pairs with a p-value of 0.05 for their correlation coefficient were considered as significantly correlated.

### Identification of actively translated transcripts

ORFquant (v1.1.0)^30^ was used to identify actively translated transcripts from the Ribo-seq alignment BAM files, which adopts a greedy approach to determine the representative transcripts and estimate the impact of RNA editing on translation. Only transcripts that were deemed translatable in both replicates were kept for subsequent analysis. Here, GENCODE M23 was used for transcript annotation.

### Enrichment analysis

ClusterProfiler (v3.14.3)^31^ was used to perform Gene Ontology (GO) biological process enrichment analysis, with an adjusted p-value of 0.05, calculated using the Benjamini-Hochberg method, being taken into account. To reduce the redundancy of enriched terms, a simplification process was implemented based on the hierarchical relationships between similar GO terms.

## RESULTS

### Characterization of high-confidence A-to-I editing sites across retina development

To obtain a global landscape of A-to-I editing in the developing mouse retina, we performed total RNA-seq to generate ten transcriptomes covering the key developmental stages including E13.5, P0, P6, P21, and P42. After quality control and data preprocessing, we applied REDItools2^22^ to characterize the RNA editing profiles. Our subsequent filtering steps, illustrated in Fig. 1A and Fig. S1A, resulted in a set of 18,778 high-confidence RNA editing sites, of which 15,269 were A-to-I editing sites, making up 81.31% of the entire set (Fig. 1B; Supplementary Table 1). This is in line with previous research findings^32,33^. Given that A-to-I editing is the most prevalent type of editing and has significant impacts on development^34–36^, we chose to focus our subsequent analysis solely on this type of editing.

**Figure 1.**
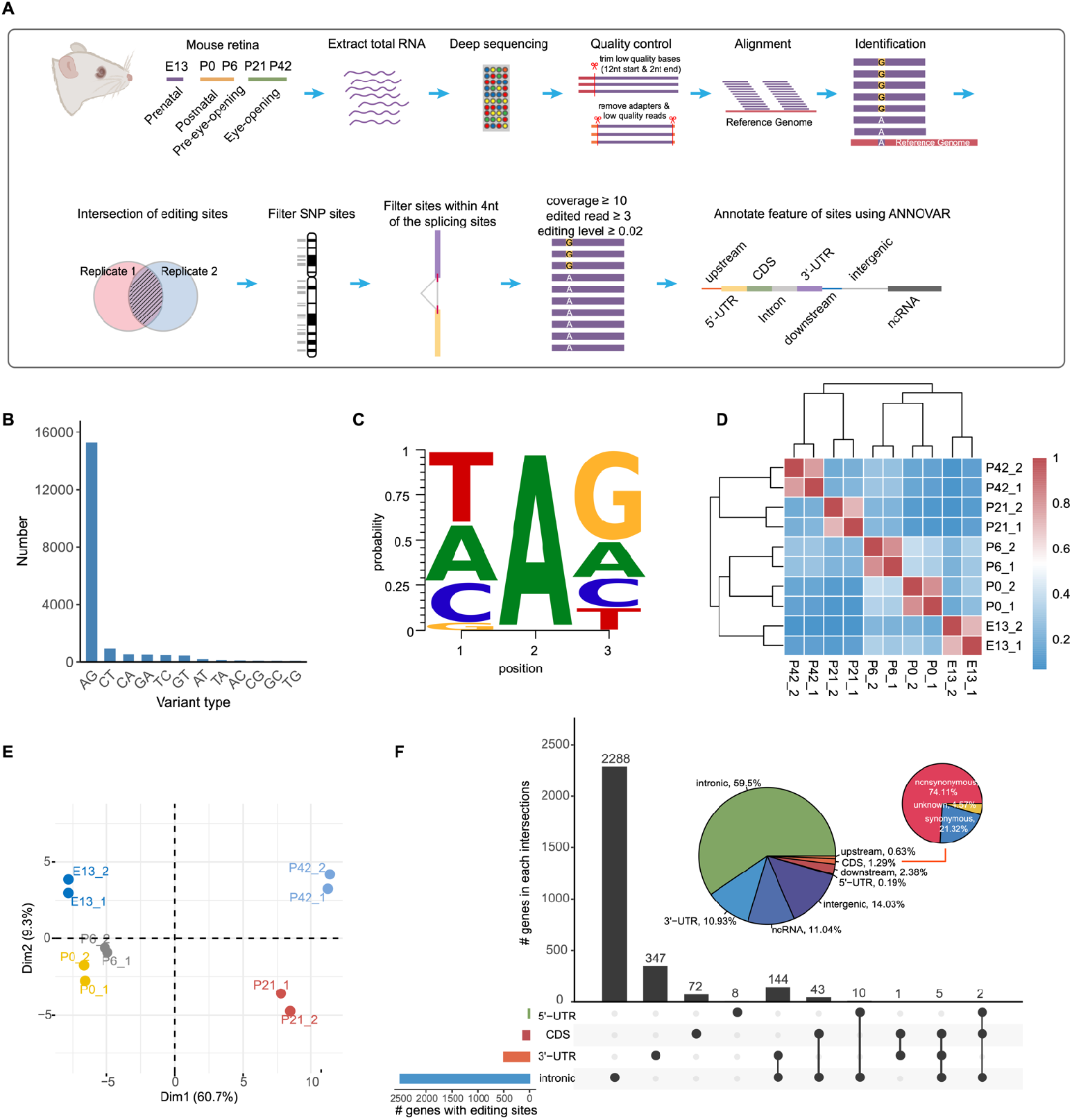
Genome-wide characterization of high-confidence A-to-I editing sites. (**A**) Schematic illustration of high-confidence A-to-I editing site identification and annotation. It summarizes the steps taken to identify and annotate A-to-I editing sites, including preprocessing of sequencing data, obtaining aligned BAM files, removing sites located in ambiguous regions and SNP sites, coverage filter, and feature annotation by ANNOVA. (**B**) Distribution of RNA editing types. Bar graph displays the number of each type of RNA editing, with ‘AG’ representing A-to-I editing sites. (**C**) Nucleotide context around A-to-I editing sites, consistent with previous findings. (**D**) Heatmap of editing levels in different developmental stages. A value closer to 1 indicates higher similarity of editing levels between samples. (**E**) Principal component analysis of editing levels in different developmental stages. (**F**) Distribution of editing sites on genomic regions. Inserted pie displays their relative fraction. Nonsynonymous refers to editing sites in the CDS that result in changes in amino acids, while synonymous refers to editing sites that do not cause such changes.

We found that the guanosine concentration was lower in the nucleotide before the editing site and higher in the nucleotide after, which aligns with the substrate requirements of ADAR editing (Fig. 1C). Hierarchical clustering analysis showed high consistency in editing levels between replicates and clear separation between developmental stages (Fig. 1D). The results of the principal component analysis reflected a developmental progression, from the embryonic (E13.5) to neonatal (P0 and P6) and then to eye-opening (P21 and P42), in line with the maturation of the retina (Fig. 1E). In agreement with previous studies^35,37,38^, we found that the majority (59.5%) of A-to-I editing sites were presented in introns, with only a small proportion (1.29%) located in coding sequences (CDS). Among those in CDS, 74.11% were non-synonymous (Fig. 1F). Overall, our analysis demonstrates the high reliability of the A-to-I editing sites we identified.

### Temporal dynamics of A-to-I editing across retina development

To investigate the temporal dynamics of A-to-I editing, we conducted an initial measurement of the number of editing sites and found a substantial variation, ranging from 325 to 11,097, among different stages of development (Fig. 2A). Despite this, a significant increase was observed over time, with a steep surge occurring after eye-opening stages. Further, we divided the editing sites into five groups based on their prevalence during development and observed that the majority (67.32%) of editing sites were exclusive to one stage, and only a negligible proportion (0.69%) of sites were shared in all stages (Fig. 2B). Analysis between different groups revealed substantial disparities in editing levels, with a trend of higher editing levels as prevalence increased (Fig. 2C). Intriguingly, shared editing sites displayed high levels of editing activity that increased as the retina developed (Fig. 2C). These sites had a preference for enrichment in 3’-UTRs (Fig. 2D). and our enrichment analysis revealed that they were often located in genes related to regulation of mRNA processing, ATP-dependent chromatin remodeling, and RNA splicing (Fig. 2E), suggesting a possible functional purpose for these sites. In contrast, D stage-specific editing sites displayed relatively low levels of editing activity, which might be due to purifying selection that impedes their editing ability or prevalence^39^.

**Figure 2.**
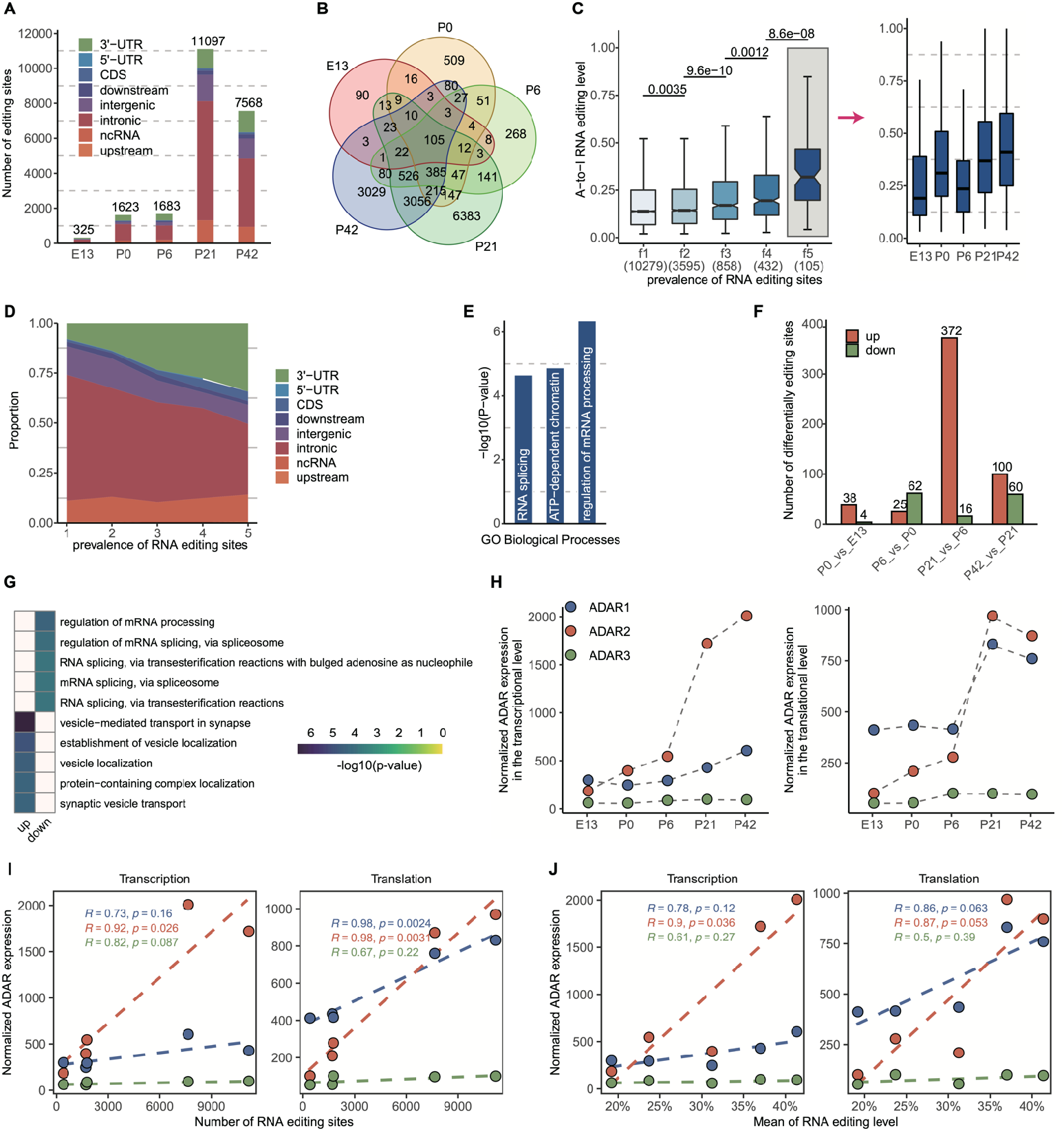
Temporal dynamics and changes of A-to-I editing. (**A**) Temporal distribution of A-to-I editing sites. (**B**) Intersection of editing sites in each stage. (**C**) Editing levels grouped by prevalence. The ‘f1’ refers to editing sties that appeared only once in the 5 stages, while ‘f5’ refers to editing sites that are shared by all 5 stages. The editing levels of f5 in different developmental stages were shown on the right. (**D**) Proportion of sites in different genomic regions with respect to prevalence. It shows the proportion of sites in 3’-UTR increasing and the proportion of sites in intronic decreasing with increasing prevalence. (**E**) Enrichment analysis of editing genes shared by all developmental stages. The top-ranked enriched GO terms are shown here. (**F**) Bar plot showing the number of differential editing sites between adjacent developmental stages. (**G**) Enriched GO biological process terms of genes with differential up- and down-regulated editing sites. (**H**) Normalized expression of three ADARs at the transcriptional (left) and translational level (right). (**I**) Pearson’s correlation analysis between editing number of all sites and ADAR expression at the transcriptional and translational levels. (**J**) Pearson’s correlation analysis between editing levels of completely-shared sites and ADAR expression at the transcriptional and translational levels, respectively.

The temporal dynamics of A-to-I editing was further studied by analyzing the differences in editing levels between adjacent developmental stages using REDITs^28^. Our results showed that a total of 625 sites underwent differential editing, with a remarkable transformation in the number of differentially edited sites before and after eye-opening (Fig. 2F and Fig. S2A; Supplementary Table 2). Our enrichment analysis indicated that as the retina developed, editing sites that experienced upregulation were primarily linked to synaptic vesicles, such as ‘vesicle-mediated transport in synapse’ and ‘synaptic vesicle transport’, while those that were downregulated were mainly related to RNA splicing, such as ‘mRNA splicing, via spliceosome’ and ‘regulation of mRNA splicing, via spliceosome’ (Fig. 2G).These findings suggest that A-to-I editing plays a crucial role in retina development, particularly in terms of modulating neurotransmitter release from synaptic vesicles and guiding alternative splicing decisions.

The expression levels of ADAR1, ADAR2, and ADAR3 play a crucial role in A-to-I editing, and therefore, we conducted further investigation into the relationship between ADAR expression and RNA editing. Our findings revealed that ADAR2 had the most noticeable increase in expression over the course of retina development, as seen at both transcriptional and translational levels, compared to the other two ADAR genes (Fig. 2H). Additionally, we found a statistically significant positive relationship between the number of editing sites and ADAR2 expression, as indicated by the Pearson’s correlation coefficients of 0.92 and 0.98 and p-values of 0.026 and 0.0031 for transcription and translation, respectively (Fig. 2I). However, no clear correlation was found between editing levels of all editing sites and ADAR expression (Fig. S3A and S3B). Despite this, a portion of the editing variability could be attributed to ADAR2 expression, as indicated by a significant positive relationship between its transcription and editing levels of completely-shared editing sites (Pearson’s r = 0.9, p-value = 0.036) and a marginal significant positive relationship between its translation and editing levels at the same sites (Pearson’s r = 0.87, p-value = 0.053) (Fig. 2J). Collectively, our results suggest that ADAR2 enzyme is crucial in controlling the developmental dynamics of the A-to-I editome.

### A-to-I editing patterns unveil functional insights on retina development

We next explored the RNA editome in greater detail to understand the changes in the editing pattern. By using mfuzz clustering^26^, we identified six distinct groups of temporal editing profiles, as shown in Fig. 3A and Supplementary Table 3. The first cluster (c1), consisting of 1,912 editing sites, showed a pattern of concurrent editing, with a sudden increase in editing levels following eye-opening. This pattern was also observed in cluster 2 (c2), which was made up of 1,968 editing sites (Fig. 3C). This editing pattern could be the result of the interaction between light and ocular tissues. Further analysis revealed that light stimulation had a positive impact on the maturation of synaptic activity in the retina following eye-opening. For instance, sites within c1 and c2 were found to be linked with genes involved in processes such as ‘regulation of long-term neuronal synaptic plasticity’, ‘endomembrane system organization’, ‘vesicle-mediated transport in synapse’ and ‘sensory perception of light stimulus’ (Fig. 3B; Supplementary Table 4). The editing patterns of clusters 3-6 were unique to their respective stages of development. Cluster 3 (c3), which was comprised of editing sites specific to P0, was characterized by functions related to those such as ‘regulation of DNA metabolic process’ ‘DNA repair’, and ‘covalent chromatin modification’, indicating that cell division and proliferation were active at this stage since this stage was marked by the formation of a substantial number of rod cells^40^. Interestingly, *Crx,* a crucial transcription for photoreceptor cell differentiation, was also edited during this stage. Cluster 4 (c4), made up of P6-specific editing sites, was characterized by functions related to those such as ‘neuron projection arborization’, and ‘negative regulation of binding’, indicating that this stage follows the production of rod cells from previous stage, reducing expression of chromatin accessibility genes such as *Nfib* to prevent excessive proliferation of rod cells and promote the wiring of neuronal connections^41,42^. Cluster 5 (c5), made up of P21-specific editing sites, was characterized by functions related to those such as ‘synapse organization’ and ‘vesicle-mediated transport in synapse’. Cluster 6 (c6), made up of P42-specific editing sites, was characterized by functions related to such as ‘covalent chromatin modification’ and ‘histone modification’, suggesting that proper editing of sites in this cluster was crucial for maintaining the functionality of the mature retina.

**Figure 3.**
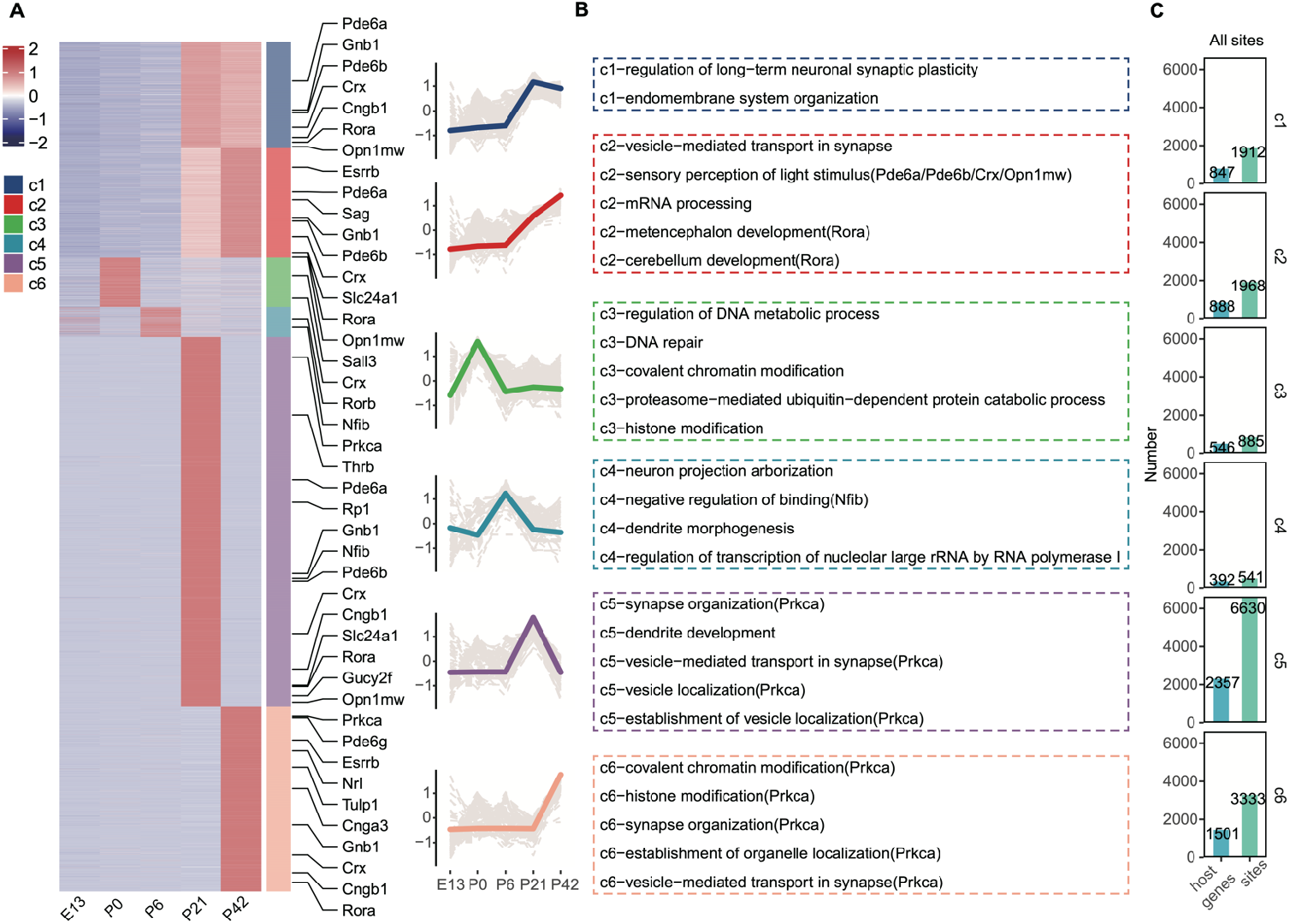
Developmental patterns of A-to-I editing. (**A**) Heatmap displaying temporal distribution of A-to-I editing during development, with the right showing the distribution of retinal markers in these patterns. The line plot depicts the editing level trends for each pattern, with colored lines indicating the normalized mean editing level of each pattern, and grey lines representing the normalized editing level of an individual editing site. (**B**) The top-ranked 5 or all enriched GO terms for each pattern in panel A are listed, with retina markers shown in parentheses after each function category. (**C**) Barplot showing the number of genes and editing sites included in each editing pattern.

### Global regulation of alternative splicing by A-to-I editing influences retina development

To gain insights into connection between RNA editing and alternative splicing, we examined their developmental interrelation. By analyzing transcriptome data (see Methods; Supplementary Table 5), we found that 68% of genes with RNA editing also exhibited alternative splicing (Fig. 4A). The presence of RNA editing was found to be significantly more prevalent among genes with splicing compared to those without, showing a strong connection between the two processes (Fig. 4B, Fisher’s exact test, p-value < 0.01). When comparing genes with and without RNA editing but both with alternative splicing, we observed that edited genes had a greater number of alternative splicing events per gene (Fig. 4C) and higher splicing efficiency (Wilcoxon rank sum test, p-value < 2.2e-16) (Fig. 4D). These results suggest that RNA editing not only can increase flexibility of splice site selection but also enhance the effectiveness of splice site recognition, thereby affecting alternative splicing decisions. Our analysis also revealed a close proximity between RNA editing and intron-retained (IR) events (Fig. S4A), suggesting that RNA editing may have a greater impact on IR events compared to other events such as EX, Alt5, Alt3, and MIC.

**Figure 4.**
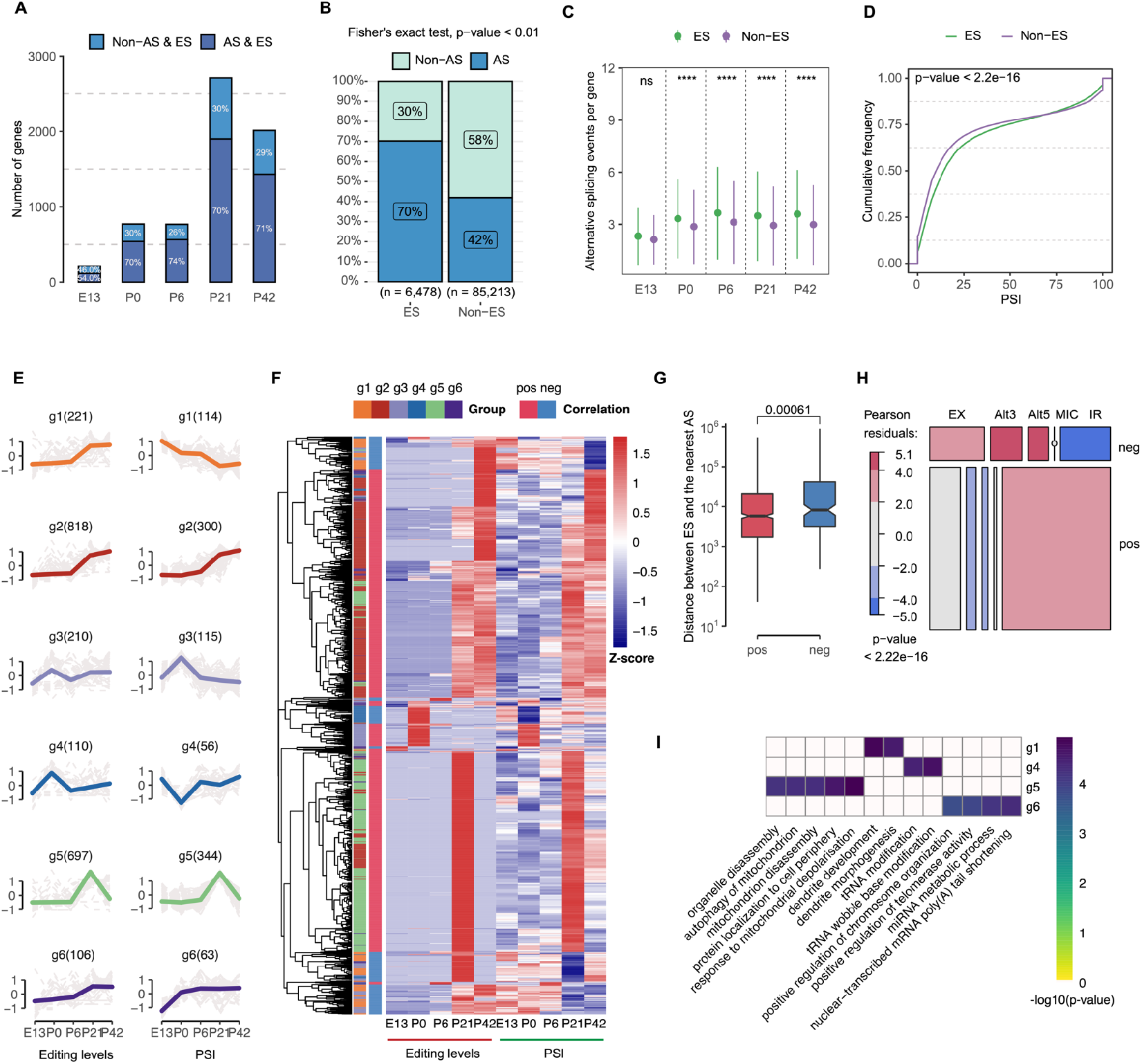
Global regulation of alternative splicing by A-to-I editing. (**A**) Proportion and number of editing genes with and without alternative splicing in each developmental stage. (**B**) Association between A-to-I RNA editing and alternative splicing, by classifying genes into four categories based on the presence or absence of editing sites and alternative splicing events, and using Fisher’s exact test to determine significance (*p*-value < 0.01). (**C**) Comparison of the average number of splicing events between genes with and without editing. (**D**) Comparison of the percent spliced-in (PSI) values between editing and non-editing genes. (**E**) Dynamic patterns of Normalized PSI values (left) and normalized editing levels (right) for strongly correlated pairs of splicing events and nearby editing sites. Colored line represents the overall trend of changes for each group, while each grey line reflects the change of a specific splicing or editing event. (**F**) Heatmap displaying the tendency of editing levels and splicing efficiency for strongly correlated pairs of splicing events and editing sites, with ‘pos’ and ‘neg’ indicating positive and negative correlation, respectively. (**G**) Comparison of the distances between paired positively correlated splicing events and editing sites with those of negatively correlated pairs. (**H**) Mosaic plot showing the distribution of different types of splicing events between positively and negatively correlated pairs of splicing events and editing sites. (**I**) Heatmap displaying the top-ranked 5 or all enriched GO terms for six strongly correlated groups.

To further examine the extent of their developmental interrelation, we focused to analyze 9,481 pairs of editing sites and their corresponding nearby splicing events. We found 2,162 pairs were strongly correlated and categorized them into six distinct groups using Mfuzz (see Methods; Figure. 4E; Supplementary Table 6). Our results showed that the changes in editing level and splicing efficiency followed a similar trajectory in groups 2, 3, 5, and 6, while groups 1 and 4 displayed a contrasting trajectory. Enrichment analysis showed that functions related to chromosome and mitochondrion, such as ‘positive regulation of chromosome organization’ and ‘mitochondrion disassembly’, were over-represented in groups 5 and 6. This suggests that editing and splicing were closely intertwined in shaping message RNA. On the other hand, functions related to development and modification, such as ‘dendrite development’ and ‘tRNA modification’, were over-represented in groups 1 and 4, indicating that splicing and editing fulfill distinct functions before and after eye-opening (Fig. 4I). Correlation analysis between editing level and splicing efficiency showed that 992 pairs of editing sites and splicing events had a significantly strong relationship (absolute Pearson’s r ≥ 0.7 and p-value ≤ 0.05) (Fig. 4F). Positively correlated editing sites and splicing events were located closer together than negatively correlated ones (Wilcoxon rank sum test, p-value = 0.00061) (Fig. 4G and Fig. S4A). IR events were more frequent than expected by chance in positive relationships, while EX events were dominant in negative relationships, indicating that the impact of RNA editing may vary depending on the type of splicing event, with high editing activity tending to favor the preservation of nearby intron or the suppression of nearby exon. (Fig. 4H). Overall, our findings suggest that A-to-I editing has a widespread effect on splicing and plays a crucial role in modulating retina development.

### Dynamic alteration of translatome plasticity conferred by A-to-I editing

Translation rate and output can be impacted by RNA editing, but the extent of this impact is yet to be determined. To shed light on this, we, we used ribosome profiling to generate translation profiles (see Methods). Our results revealed that the combination of RNA editing and splicing (AS & ES) resulted in the highest average number of actively translated transcripts per gene. This was followed by the group with only splicing (AS & Non-ES), then by the group with only editing (Non-AS & ES), and finally by the group with neither editing nor splicing (Non-AS & Non-ES) (Fig. 5A). However, the Non-AS & Non-ES group had the highest translational efficiency, followed by the Non-AS & ES, AS & Non-ES, and AS & ES groups (Fig. 5B). These findings indicate that editing and splicing can increase coding capacity and diversify the translatome, with a synergistic effect when used together, for example, the editing level and splicing efficiency of retina-specific gene Pcdh15, Stx3, Pde6b, and Tia1 synergistically inhibit their TE (Fig. S5A). Notably, the rise in translatome diversity was accompanied by usage increase in RNA editing and splicing, with the latter having a more pronounced impact on translational efficiency of gene than the former.

**Figure 5.**
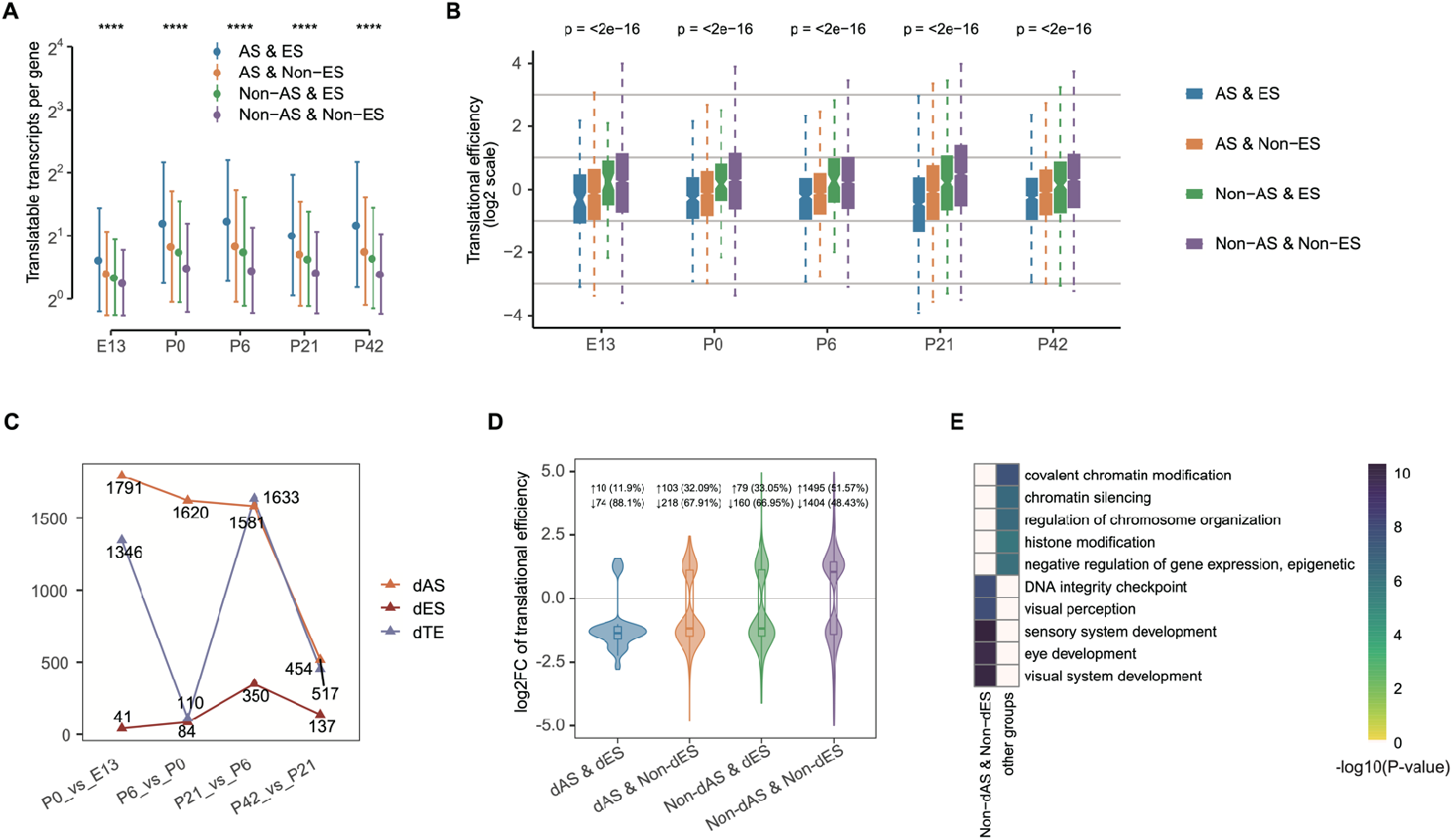
A-to-I editing induces dynamic changes in translatome plasticity. (**A**) Point-range plot displaying the mean number of translatable transcripts for four groups of genes classified based on the presence or absence of splicing events or editing sites. (**B**) Boxplot showing translational efficiency between different groups of genes. (**C**) Number of genes with differential splicing events, differential editing sites, and differential translational efficiency in pairwise comparisons. (**D**) Violin plot comparing translational efficiency among four gene groups classified based on whether they exhibited differential editing or differential splicing efficiency. (**E**) Heatmap showing the top-ranked 5 enriched biological process GO terms for different gene groups in panel D.

In light of these findings, we further examine the effect of RNA editing on the translatome dynamicity. We performed differential translational efficiency (dTE) analysis between adjacent developmental stages and found 2,936 dTE genes that mirrored retina development, with two pronounced peaks in gene number between E13 and P0, and P6 and P21 (see Methods and Fig. 5C; Supplementary Table 7). In parallel, we also found 3,453 differential splicing efficiency (dPSI) genes and 457 differential editing (dEL) genes (see Methods). When dTE genes were classified into four groups based on their dPSI or dEL status, we observed that the Non-dAS & Non-dES group had a close balance between up- and down-regulated dTE genes, with 51.57% and 48.43%, respectively. The balance was disrupted in the presence upon of dPSI or dEL, resulting in the majority of dTE genes being down-regulated in the Non-dAS & dES (66.95%), dAS & Non-dES (67.91%), and dAS & dES (88.1%) groups (Fig. 5D). These results indicate that RNA editing and splicing can serve as a buffering mechanism to reduce translational efficiency, with both having a coordinated effect. Enrichment analysis further revealed that only the Non-dAS & Non-dES group had an over-representation of functions related to retina development, such as ‘lens development in camera-type eye’, ‘visual perception’, and ‘visual system development’, while the other three groups had an over-representation of functions related to the basic processes of life, such as ‘chromatin silencing’, ‘RNA splicing’, ‘‘’nuclear export’, and ‘regulation of chromosome organization’ (Fig. 5E). These findings suggest that RNA editing and splicing play an important role in preserving multiple basic cellular processes.

## DISCUSSION

Adar-mediated A-to-I editing has been established as crucial for normal development of organisms^33,43,44^. Disruptions to Adar can lead to serious consequences such as locomotion and neuron defects seen in flies with mutant Adar^45^. However, the contribution of A-to-I editing to retina development has yet to be fully understood. We herein sought to understand the role of A-to-I editing in mouse retina development, with a specific focus on its regulatory effect on gene translation. By analyzing tens of thousands of editing events in mouse retina development, we created a detailed temporal map of the A-to-I editome. Our findings provide novel insights into functional impact of A-to-I editing on retina development and highlight its significance in regulating alternative splicing to dynamically control the retina translatome.

A-to-I editing is catalyzed by ADARs, of which there are three members in mammals: ADAR1, ADAR2, and ADAR3. Our results suggest that ADAR2, and not ADAR1 or ADAR3, is primarily responsible for dynamic changes in A-to-I editing, underlining the intricate regulatory role of ADAR2 in retina development. The editing patterns produced by temporal changes in ADAR2 display specific and continuous characteristics, with the majority of editing sites exhibiting stage-specific changes during development, presumably in order to fulfil unique requirements of specialized retinal functions. Our findings indicate that retina development is tightly regulated by A-to-I editing in specific and continuous manners to ensure the proper generation of cell types and the formation of functional neuronal circuitry. Altogether, these results demonstrate the important role of A-to-I editing in retina development.

The frequent observation of splicing in functions that were significantly enriched was noteworthy. This confirmed the close relationship between RNA editing and splicing, which was in agreement with prior findings that have emphasized the interplay between the two^14^. Our examination of alternative splicing events further showed that A-to-I editing had the ability to increase the number of splicing events by providing potential donor or acceptor sites, and that it could affect splicing efficiency by changing the structure and stability of sequences. It is clear that A-to-I editing and alternative splicing are deeply interconnected, with the former having a profound impact on the latter. Notably, RNA editing and splicing are both key pre-mRNA processing steps that can introduce substantial modifications to final gene products^14,15^. Although the ability to dynamically regulate transcriptome diversity has been established, the potential influence of RNA editing and splicing on translatome remains poorly understood. Our results indicate that A-to-I editing and splicing play crucial roles in enhancing translatome diversity and are often utilized to modify translatome dynamics. Specifically, A-to-I editing was found to have the potential to decrease translational efficiency through interaction with splicing. To our knowledge, this study offers a pioneering depiction of the complex interplay between RNA editing, alternative splicing, and translation.

The current study provides substantial functional predictions and in silico confirmation, however, its limitations include lack of experimental validation. The identification of RNA editing sites is a challenging task. Although the screening process was designed to guarantee the accuracy of the sites, additional confirmation through Sanger Sequencing could further advance our exploration into their biological significance. Future studies focused on uncovering the relationship between RNA editing enzymes and splicing machineries will deepen our knowledge of retina development mechanisms. Furthermore, including a broader range of time points in the analysis, beyond the current focus on major phases of retina development, will provide a more comprehensive understanding of RNA editing and its role in neurological condition. Integrating with other technologies such as scRibo-seq will allow for a more in-depth analysis of RNA editing’s impact on cell type-specific translation and regulation.

## DATA AVAILABILITY

The total RNA-seq and Ribo-seq sequencing data is available in the NCBI Gene Expression Omnibus (GEO) with the accession number GSE104884.

## SUPPLEMENTARY MATERIALS

Supplementary materials can be found online.

## ACKNOWLEDGEMENT

We thank the support for the Center for Precision Medicine, Sun Yat-sen University.

## AUTHOR CONTRIBUTIONS

Z.X. and H.W.W. supervised the project; J.Q.Y. performed RNA-seq and Ribo-seq library construction; L.D.Y., L.Y., and R.Z. analyzed and interpreted the data; L.D.Y., H.W.W., and Z.X. wrote the manuscript; All authors approved the manuscript.

## CONFLICTS OF INTEREST

The authors have no conflicts of interest to declare.

## FUNDING

This work was supported by the National Key R&D Program of China [2019YFA0904400 to Z.X.], National Natural Science Foundation of China [32270700 to H.W.W], and Guangzhou Science and Technology Project (202201020336 to Z.X.).

## Notes

### Competing Interest Statement

The authors have declared no competing interest.

